# Deep Mutational Scan of a cardiac sodium channel voltage sensor

**DOI:** 10.1101/773887

**Authors:** Andrew M. Glazer, Brett M. Kroncke, Kenneth A. Matreyek, Tao Yang, Yuko Wada, Tiffany Shields, Joe-Elie Salem, Douglas M. Fowler, Dan M. Roden

## Abstract

Variants in ion channel genes have classically been studied in low-throughput by patch clamping. Deep Mutational Scanning (DMS) is a complementary approach that can simultaneously assess function of thousands of variants. We have developed and validated a method to perform a DMS of variants in *SCN5A*, which encodes the major voltage-gated sodium channel in the heart. We created a library of nearly all possible variants in a 36 base region of *SCN5A* in the S4 voltage sensor of domain IV and stably integrated the library into HEK293T cells. In preliminary experiments, challenge with three drugs (veratridine, brevetoxin, and ouabain) could discriminate wildtype channels from gain and loss of function pathogenic variants. High-throughput sequencing of the pre- and post-drug challenge pools was used to count the prevalence of each variant and identify variants with abnormal function. The DMS scores identified 40 putative gain of function and 33 putative loss of function variants. For 8/9 variants, patch clamping data was consistent with the scores. These experiments demonstrate the accuracy of a high-throughput *in vitro* scan of *SCN5A* variant function, which can be used to identify deleterious variants in *SCN5A* and other ion channel genes.

## Introduction

*SCN5A* encodes the major voltage-gated sodium channel in the heart, Na_V_1.5. This protein consists of 2016 residues with 24 membrane-spanning helices in four homologous domains (Figure 1). Variants in *SCN5A* cause multiple distinct genetic arrhythmia syndromes: complete or partial loss of function variants are the most common genetic cause of Brugada Syndrome (BrS1)^1^ and gain of function variants are the third most common genetic cause of congenital Long QT Syndrome (LQT3; Figure 1).^2^ *SCN5A* variants have also been associated with isolated conduction system disease, atrial fibrillation, and dilated cardiomyopathy.^3^ BrS1 and LQT3 can cause sudden cardiac death, and treatment in diagnosed subjects can prevent this lethal outcome.^4^ The function of *SCN5A* variants is conventionally assayed in patch clamp experiments which are highly predictive of *SCN5A* variant disease risk (Figure 1A).^5^ In particular, *SCN5A* variants that display a reduction in peak current (“loss of function”) are associated with BrS1,^5, 6^ and variants that disrupt fast inactivation and increase late current (“gain of function”) are associated with LQT3 (Figure 1B).^5, 7^

**Figure 1:**
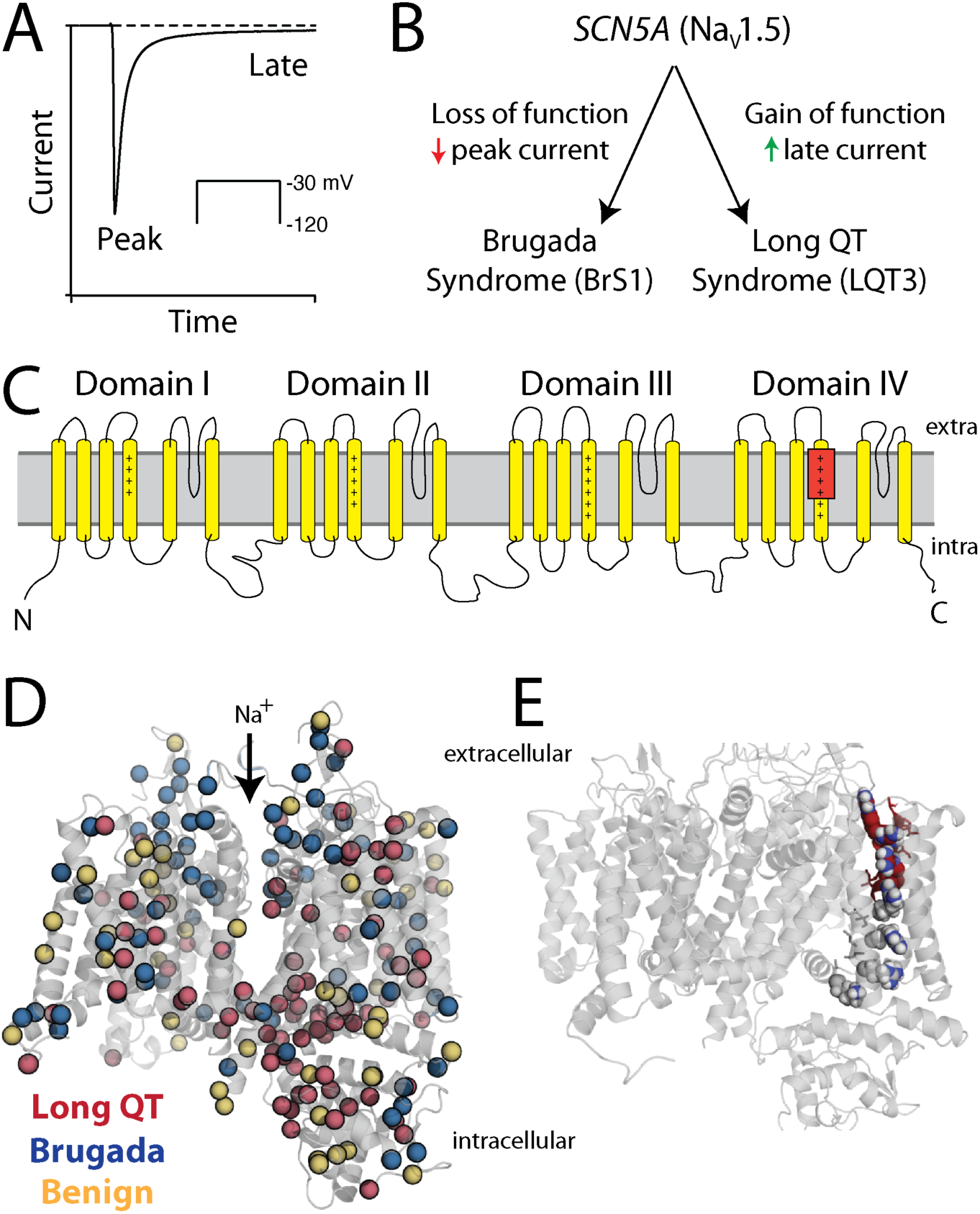
Diagram of Na_V_1.5. A) Example Na_V_1.5 patch clamp electrophysiology result. Peak and late currents are indicated. B) Diagram of the major *SCN5A*/Na_V_1.5 associations with disease. Additional minor arrhythmia associations are not shown. C) Schematic of Na_V_1.5 structure. The red box indicates the 12 amino acid region (#1621-1632) used for the Deep Mutational Scan. Positively charged residues (mostly arginines) in the S4 helices are indicated with “+”. D) Model of Na_V_1.5 structure built from a homologous CryoEM structure.^52^ Residues with >20% estimated penetrance of Long QT or Brugada Syndrome are indicated in red/blue, respectively, and residues with <20% estimated penetrance for either disease are indicated in yellow.^5^ As discussed in the text, a small subset of variants show both BrS1 and LQT3 pathology. E) Model of Na_V_1.5 structure with 12 amino acid target region in red. Arginine residues are shown in blue (4 of which lie within the 12 aa target region).

Because BrS1 and LQT3 are treatable disorders, the American College of Medical Genetics and Genomics (ACMG) has identified *SCN5A* as one of 59 “actionable” genes for which incidental pathogenic variants should be reported to the clinician.^8, 9^ However, our group has identified a surprisingly high degree of disagreement over which *SCN5A* variants are considered pathogenic.^10^ Two major ACMG classification criteria rely on functional data: “PS3” (Pathogenic Strong #3) and “BS3” (Benign Strong #3).^8, 11^ “Well established functional studies supportive of a damaging effect on the gene or gene product” can be used to achieve the PS3 criterion and “well-established functional studies that show no deleterious effect” can be used to achieve the BS3 criterion.^11^ Unfortunately, Variants of Uncertain Significance (VUS) currently outnumber all other categories in the ClinVar database and their number is rapidly growing.^12^ For *SCN5A,* 698 variants are listed as VUS or have conflicting interpretations in ClinVar,^13^ and a total of 1,712 *SCN5A* variants have been observed in at least one individual to date in the literature or in the gnomAD database, a majority of which are classified as VUS’s.^5, 14^ As genomic medicine becomes commonplace, the problem of variant classification will continue to grow unless accurate, high-throughput variant annotation strategies are developed.

Deep mutational scanning (DMS) is a massively parallel method for evaluating variant effects. In a DMS, a pool of variants is created, and a selection for the gene product’s activity is applied. Each variant in the pre- and post-selection pools is counted using high-throughput DNA sequencing.^15^ While DMS has been successfully implemented for many genes and protein domains,^16–19^ the approach has not been adapted to arrhythmia-associated ion channel genes. In this work, we demonstrate a method to apply DMS to *SCN5A* to identify both loss and gain of function variants, enabling multiplex variant assessment in this important disease gene. We measure the effect of 248 variants in the voltage sensor (S4) segment of transmembrane domain IV, a hotspot region known to harbor multiple disease-associated gain and loss of function variants.^20–23^ We identify 40 putative gain of function and 33 putative loss of function variants, 8/9 of which were validated by patch clamp data.

## Results

### A triple drug assay can identify *SCN5A* loss and gain of function variants

In order to characterize many *SCN5A* variants simultaneously, we attempted to develop a parallelizable *in vitro* assay of Na_V_1.5 function. We adapted and optimized a previously described HEK293T cell toxicity assay used for screening environmental samples for sodium channel-modulating toxins.^24^ The assay uses two sodium channel agonists, veratridine and brevetoxin, and the Na+/K+ exchanger inhibitor ouabain. The Na_V_1.5 agonists synergistically bias the channel towards an open state, resulting in sodium influx.^25^ Ouabain causes increased sensitivity to sodium overload, resulting in death of cells expressing functional sodium channels on their surface.^26^ To measure the effects of the triple drug assay, we used flow cytometry to quantify GFP+ percentages in cells transiently transfected with a p*SCN5A*-IRES-GFP plasmid. We determined optimal drug concentrations (10 µM ouabain, 25 µM veratridine, and 250 ng/ml brevetoxin) and treatment time (5 hours) to maximize selection against expressing a high level of wildtype *SCN5A* (Figure 2A-B, S1). When all three drugs were added at the final dose, we observed a strong selection (35% normalized cell survival rate) against cells expressing wildtype *SCN5A* (Figure 2B).

**Figure 2:**
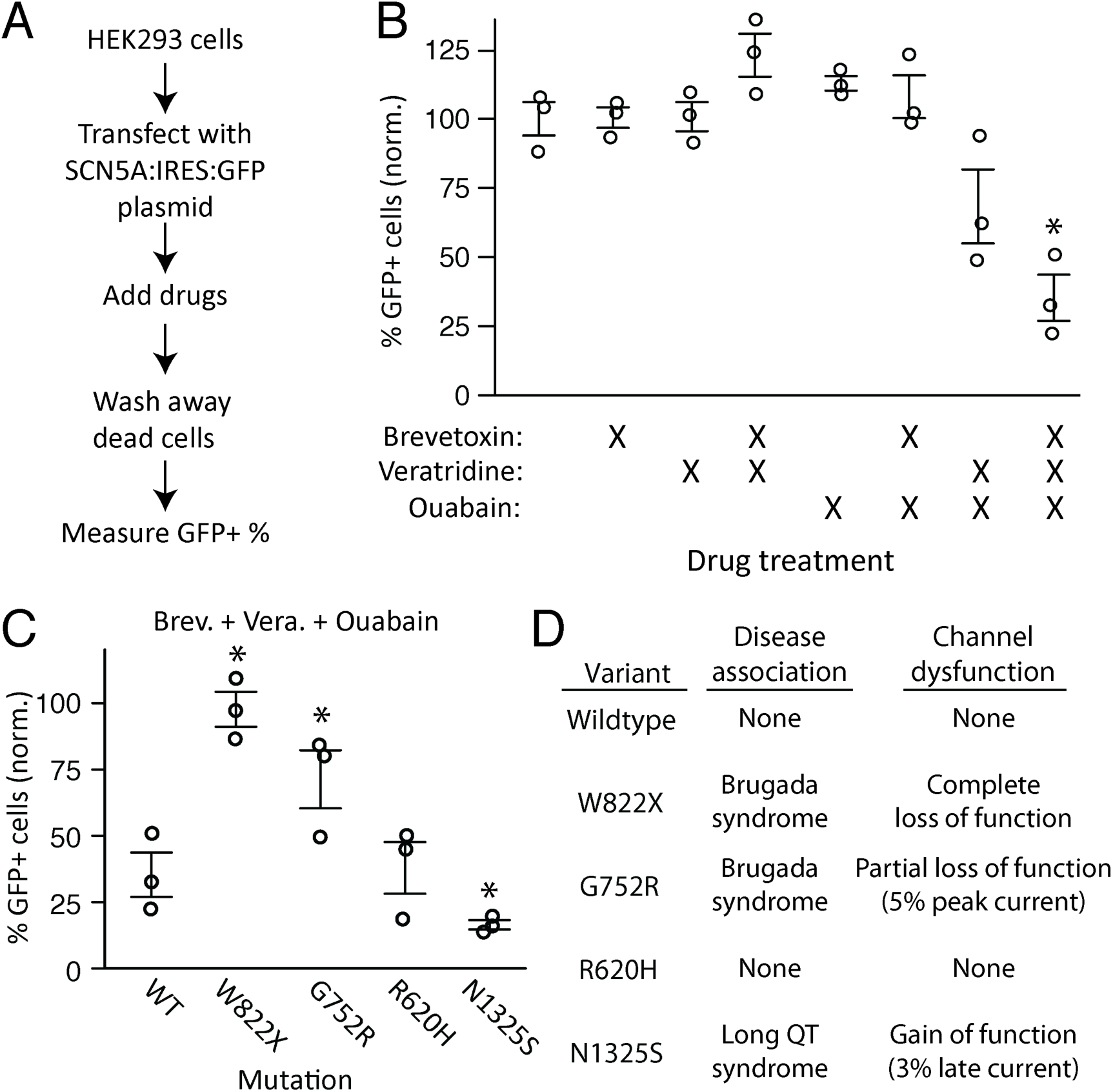
Triple drug treatment distinguishes between wildtype, gain of function, and loss of function *SCN5A* mutations. A) Schematic of method to detect selection against *SCN5A* variants after drug treatment. B) HEK293T cells were transfected with wildtype *SCN5A* and treated with the 8 possible combinations of veratridine, brevetoxin, and ouabain. Experiments were done in triplicate, and error bars indicate standard error of the mean. C) HEK293T cells were transfected with previously studied *SCN5A* variants and treated with veratridine, brevetoxin, and ouabain (“triple drug treatment”). Percentage GFP+ cells (normalized) indicates the rate of cell survival after drug challenge as measured by percentage of cells expressing GFP, normalized to a no-drug control. Three replicates were analyzed for each variant or condition. Error bars indicate standard error of the mean. *p<0.05, two-tailed t-test compared to no-drug control (B) or wildtype (C). D) Previously published disease associations and functional electrophysiologic characterizations.^27–30^

We next tested the triple drug treatment against an allelic series of four previously described *SCN5A* variants: 2 BrS1-associated loss of function variants, 1 LQT3-associated gain of function variant, and a non-synonymous variant that does not alter channel function.^27–30^ When cells were treated with the triple drug mixture, we observed a graded response of cell survival depending on the degree and direction of channel dysfunction (Figure 2C-D). Specifically, HEK293T cells expressing wildtype *SCN5A* had 35 +/− 8% normalized GFP+ cells (mean +/− S.E.M.). This value was similar to a non-disruptive mutation (R620H, 38 +/− 10%). Cells expressing a gain of function mutation (N1325S, 16 +/− 2%) had decreased viability compared to wildtype. Cells expressing a complete loss of function mutation (W822X, 98 +/− 7%) or partial loss of function mutation (G752R, 71+/− 11%) had increased viability relative to wildtype. All tested variants except R620H had significantly altered GFP+ cell survival rates compared to wildtype (p<0.05, n=3 replicates/variant). These results indicate that the triple drug assay can distinguish loss of function and gain of function *SCN5A* variants from wildtype.

### A Deep Mutational Scan of a 12 amino acid region of *SCN5A*

We next performed a Deep Mutational Scan of *SCN5A* by applying the triple drug treatment to a library of *SCN5A* variants (Figure 3A). We chose a 12 amino acid region in the voltage-sensing helix 4 of transmembrane domain IV (Figure 1C) and performed comprehensive mutagenesis on this region using inverse PCR with degenerate primers (Figure S2).^31^ We inserted an 18-mer random barcode into the plasmids (Figure S3). Subassembly to identify barcode-mutation relationships identified 7,205 unique barcoded plasmids, including all possible nonsense and synonymous amino acid changes, and 224 of the 228 possible nonsynonymous changes (Figure S4, Table S1). This library of mutant plasmids was integrated into HEK TetBxb1BFP cells such that each cell expressed only one *SCN5A* variant.^32^ Single integrant cells, which became mCherry+, BFP-following plasmid integration, were isolated using fluorescence activated cell sorting. After cell recovery from sorting, a pre-drug sample was collected. Then, the triple drug treatment was applied, dead cells were removed by repeated media changes, and a post-drug sample was collected. For both pre- and post-drug samples, barcodes were sequenced on an Illumina MiSeq sequencer. Six independent replicates were collected, and data analyzed as described below.

**Figure 3.**
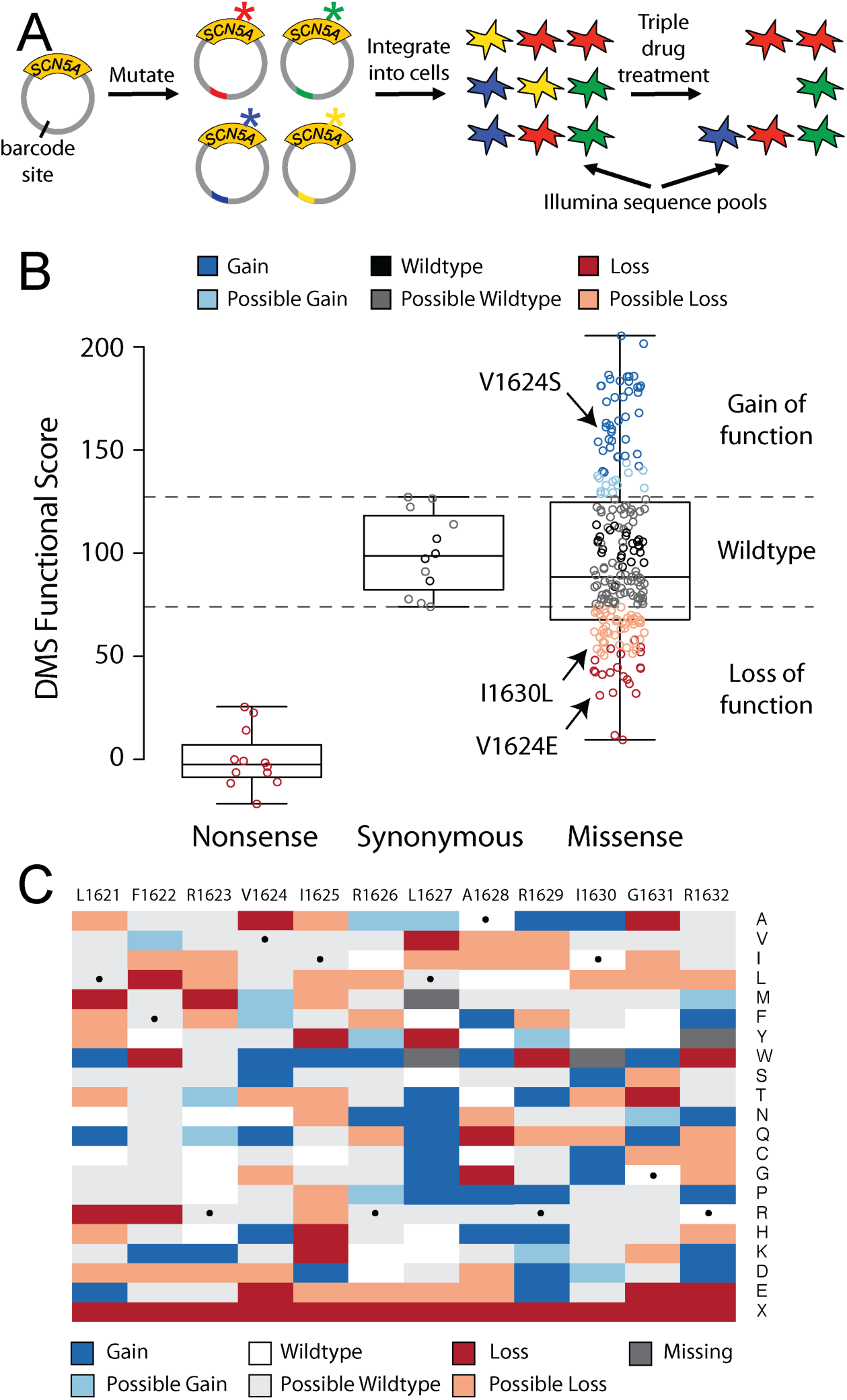
Deep Mutational Scan accurately identifies *SCN5A* variants with normal and abnormal function. A) Diagram of *SCN5A* Deep Mutational Scan approach. NNN-mutagenic primers were used to generate nearly all possible variants; each plasmid was also tagged with a unique barcode. The pool of all plasmids was then transfected into HEK cells engineered to include a “landing pad” that accommodates only a single integrant per cell. The pool was then challenged with three drugs (ouabain, veratridine, and brevetoxin) to bias survival toward cells that expressed functional protein. High-throughput sequencing of the barcodes was then used to calculate a functional score. B) Boxplot of Deep Mutational Scan scores. Each data point is a variant. Deep Mutational Scan scores were averaged across 6 replicates and calculated as the log_10_(prevalence initial library/prevalence post-drug). The scores were linearly scaled so that the mean nonsense variant had a score of 0 and the mean synonymous variant had a score of 100. C) Heatmap of Deep Mutational Scan scores. Scores were calculated as described in B. Synonymous variants are indicated with a black dot.

### Deep Mutational Scan identifies loss and gain of function Na_V_1.5 variants

To determine if the Deep Mutational Scan successfully applied a negative selection against sodium channel function, we examined the fold changes of mutants after selection by functional class. Relative to their prevalence in the initial library, the nonsense variants were enriched by 2.2 +/− 1.2-fold in the post-drug pool (mean +/− S.E.M.), whereas synonymous variants and nonsynonymous variants were depleted by 25.9 +/− 1.3-fold and 24.8 +/− 1.1-fold, respectively (Figure S5). For each variant, we calculated a DMS score from the log_10_ ratio of post-drug to initial library prevalence, linearly normalized so that the mean synonymous score was 100 and the mean nonsense score was 0. Four variants absent from the initial plasmid library were excluded. As a result, we generated DMS scores for 224 of the 228 possible nonsynonymous variants and all 24 possible nonsense/synonymous variants. To determine experimental variability, we calculated normalized DMS scores from each of the six replicates individually and showed that replicate scores correlated (mean Pearson’s r=0.63 and Spearman’s r=0.59).

To increase the accuracy of the scores, mean DMS scores were calculated by averaging the scores from all six replicates, and 95% confidence intervals were calculated from the replicate scores. The range of the 12 synonymous scores was 74 to 127 and the range of the 12 nonsense scores was −22 to 26 (p<0.001). Variants with mean and 95% confidence intervals higher or lower than the synonymous range were classified as gain or loss of function variants, respectively (Figure 3B). Variants with means outside the synonymous range but 95% confidence intervals overlapping the range were classified as possibly gain or loss of function. Using these classifications, 40 gain of function, 14 possibly gain of function, 33 loss of function, and 51 possibly loss of function variants were identified (Figure 3B). All normalized DMS scores, standard errors, 95% confidence intervals, and interpretations (e.g. gain/loss of function) are presented in File S1.

A characteristic feature of ion channel S4 helix voltage sensors is the presence of positively charged residues (mainly arginines in the case of Na_V_1.5) at every third position.^33^ These residues are important for voltage sensing and voltage-dependent activation, and domain IV arginines are also linked to channel inactivation.^34–36^ The first four S4 arginines were included in the 12 amino acid region (R1623, R1626, R1629, and R1632; Figure 1C). In the literature, 3 gain of function variants have been identified in R1623 or R1626 and 3 loss of function variants in R1629 and R1632 (Table 1).^20–23, 37, 38^ The three previously-studied partial loss of function BrS1 variants all had low Deep Mutational Scan scores (62, 66, and 69). Two of three previously studied gain of function LQT3 variants had elevated Deep Mutational Scan scores (129 and 144), while a third mild gain of function variant had a near-wildtype score (95). Overall, we observed a complex pattern of gain and loss of function scores in the arginine residues (Figure 3C). Each arginine reside had at least 1 gain of function variant and 1 loss of function variant. We also observed a high prevalence of gain (n=26) and loss of function (n=17) nonsynonymous variants in the 8 non-arginine residues in the target region.

**Table 1:** Deep Mutational Scan scores correlate with patch clamp dysfunction and disease association.

| **Variant** | **DMS Score** | **Patch clamp result** | **Disease association** | **Controls** | **LQT patients** | **Brugada patients** |
| --- | --- | --- | --- | --- | --- | --- |
| R1623Q | 129.5 | Gain<sup>20</sup> | Long QT<sup>46</sup> | 0 | 15 | 0 |
| R1626P | 143.9 | Gain<sup>22</sup> | Long QT<sup>22</sup> | 0 | 2 | 0 |
| R1626H | 95.4 | Mild Gain<sup>38</sup> | Long QT(?)<sup>47</sup> | 11 | 2 | 0 |
| R1629Q | 69.1 | Partial Loss<sup>23</sup> | Brugada<sup>23</sup> | 5 | 0 | 4 |
| R1632C | 61.9 | Partial Loss<sup>21</sup> | Brugada<sup>21</sup> | 1 | 0 | 2 |
| R1632H | 66.6 | Partial Loss<sup>37</sup> | Brugada<sup>53</sup> | 3 | 0 | 10 |
| V1624S | 154.1 | Gain | N/A | 0 | 0 | 0 |
| I1630L | 59.2 | Partial Loss | N/A | 0 | 0 | 0 |
| V1624E | 32.4 | Loss | N/A | 0 | 0 | 0 |

\*Patient counts are taken from a curation of the *SCN5A* variant literature.<sup>5</sup> Patch clamp data are from the literature or this study (Figure 4, Table S2). For patient counts a representative reference was chosen.

### Patch clamp experiments validate Deep Mutational Scan scores

In order to further validate the Deep Mutational Scan scores, we compared the scores to gold-standard patch clamp measurements in HEK293T cells. As mentioned above, 5/6 previously studied nonsynonymous variants in arginine residues in the target region had DMS scores consistent with the previous patch clamp results (Table 1). In addition to these six previously studied variants, we studied three other variants, predicted to have loss or gain of function phenotypes based on their Deep Mutational Scan scores and not previously investigated by patch clamp. All three variants had the predicted functional change when characterized by the patch clamp (Figure 4, Table S2). V1624E (score 32) had no detectable current, I1630L (score 59) had 29.7% of the wildtype level of peak current, and V1624S (score 154) had a slight reduction in peak current (76.6% of wildtype) but a large increase in late current (1.95% of peak) compared to wildtype (0.29% of peak). These data show that the Deep Mutational Scan successfully identified loss and gain of function *SCN5A* variants.

**Figure 4.**
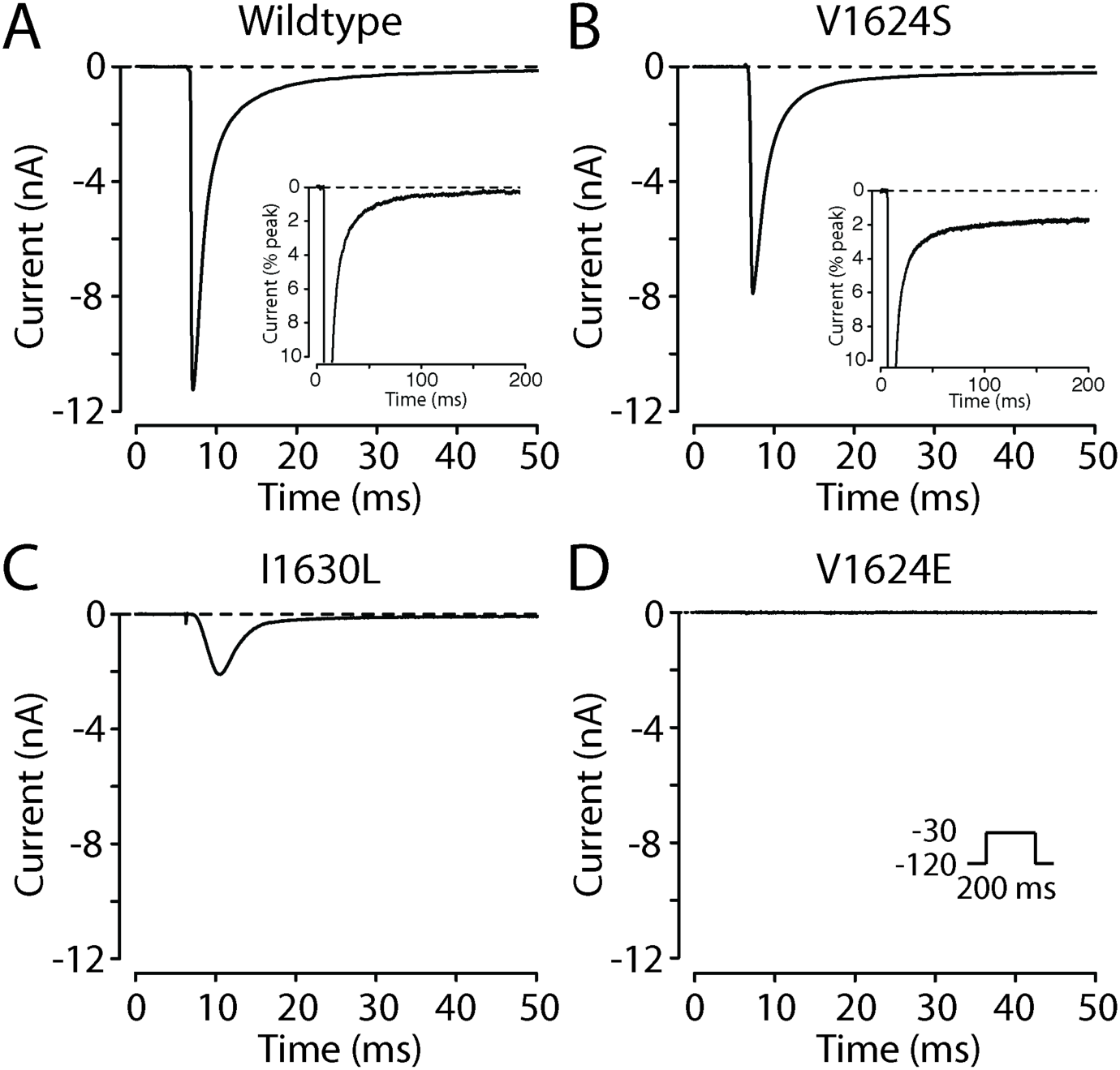
*SCN5A* Deep Mutational Scan accurately predicts mutational effects. A-D) Voltage clamp of wildtype and mutant *SCN5A*. Cells were exposed to a 200 ms depolarizing pulse from a holding potential of −120 mV to −30 mV. A) WT, B) V1624S (DMS predicted gain of function), C) I1630L (DMS predicted partial loss of function), D) V1624E (DMS predicted loss of function). A, B) Insets show increased late current in V1624S compared to WT. Further patch clamp data and statistics are presented in Table S2.

## Discussion

Deep Mutational Scans have the advantage of making functional measurements of variants at a scale (hundreds to thousands of variants) that is able to keep pace with the accelerating rate of variants being found in sequencing projects.^12^ Importantly, the Deep Mutational Scan approach presented here can generate functional scores in a multiplexed manner. Therefore, in terms of scale and cost-effectiveness, the Deep Mutational Scan approach compares favorably with both conventional manual patch-clamp as well as emerging methods such as automated patch-clamp or optical screening in multiwell plates.^39^

A challenge of assaying Na_V_1.5 function is that the channels’ default state is closed, and they open for only ∼5 ms in response to a rapid increase in the cell’s membrane potential before inactivating. In this study, we used two sodium channel agonists to bias the cell towards the open state and promote sodium influx, which we made more toxic to the cell by treatment with an inhibitor of the Na+/K+ exchanger. Simultaneous treatment with both classes of drug was required for negative selection, and the highest selection was observed with two channel agonists (Figure 2), consistent with previous findings that veratridine and brevetoxin act in a synergistic manner.^25, 40^ Channel agonists are commonly used in other high-throughput screening approaches,^41, 42^ and therefore might be used to help develop analogous functional assays for other ion channels.

The *SCN5A* Deep Mutational Scan identified both loss and gain of function variants, which is especially useful for *SCN5A* because both classes of variants are associated with clinical arrhythmia syndromes. The accuracy of these scores is demonstrated by the 1) the correlation of replicate measurements, 2) the strong difference in scores between nonsense and synonymous variants, and 3) the consistency between the scores and channel dysfunctions for 8/9 variants as measured by patch clamping.

The positively charged residues (e.g. arginines) of the four S4 helices are classically considered to implement voltage sensing.^36^ The domain IV S4 helix, however, is less tightly coupled to channel activation and is linked to channel inactivation.^34, 35, 43^ Na_V_1.5 variants in this helix can cause gain or loss of channel function by affecting the inactivation process.^20–23, 37, 38^ For example, R1629Q has wildtype-like peak current, but causes a leftward shift in steady-state inactivation and an increased inactivation rate.^23^ Our DMS assay suggests an overall partial loss of function character that is consistent with these data and the clinical manifestation of Brugada Syndrome^23, 44^ (Table 1). By contrast, R1623Q has wildtype-like peak current and a small leftward shift in steady state inactivation, but a large decrease in inactivation rate.^20, 45^ Our DMS assay suggests an overall gain of function character, consistent with these data and the clinical manifestation of Long QT Syndrome^46, 47^ (Table 1). These results suggest that our assay is sensitive to a variety of disease-relevant perturbations, in addition to canonical classes of variants like increased late current (e.g. V1624S) and decreased peak current (e.g. V1624E, I1630L).

In addition to previously studied variants, this work provides an expanded picture of the impact of nonsynonymous variants in this helix. We observed a complex pattern of mutational effects at the arginine residues, with at least one gain and loss of function variant at each position. We also observed a high rate of gain and loss of function variants in non-arginine residues in this region, 3/3 of which were validated by patch clamping. This complexity highlights the difficulty of predicting variant effects and illustrates the value of empirically determining *SCN5A* variant function/dysfunction. Traditional methods such as patch clamping are critical but have investigated only 219 of the 1,390 *SCN5A* missense variants that have been observed in at least one individual to date.^5, 7^ Our Deep Mutational Scan approach has the potential to gather functional data at scale to help assess the growing number of observed rare *SCN5A* variants.

### Limitations

The DMS appears appears to successfully distinguish multiple classes of disease-relevant channel dysfunction. However, there may be certain types of dysfunction (e.g. complex or subtle gating alterations) that are not captured by this approach. Several *SCN5A* “overlap” mutants have been identified that have both partial loss-of-function (decreased surface abundance) and gain-of-function (large late current) features;^48^ these variants may falsely appear as wildtype-like in the Deep Mutational Scan. Variants that disrupt binding of veratridine or brevetoxin to Na_V_1.5 may be falsely interpreted as loss of function. DMS scores for residues at these drug binding sites, which have been mapped and do not include the region in this study,^49^ should be excluded.

### Conclusions

We present a high-throughput approach that can scan for loss and gain of function variants in the cardiac sodium channel *SCN5A*. This method can be used to identify variants associated with Brugada or Long QT syndromes. This approach may be adapted for other ion channels implicated in Mendelian disease to help improve assessments of variant disease risk.

## Methods

### Mutagenesis

Single point mutations were introduced into *SCN5A* using the QuikChange Lightning kit (Agilent) and verified by Sanger sequencing. Comprehensive codon mutagenesis was performed using an inverse PCR method.^31^ We performed inverse PCR on a plasmid containing a portion (AA#1533-1785) of the wild-type *SCN5A* gene using one primer pair per codon (diagrams of the mutagenesis and cloning methods are provided in Figures S2 and S3, respectively). The first three nucleotides of the forward primer were “NNN” (each nucleotide is a mix of A/T/G/C). PCR products corresponding to the 12 mutated codons were pooled and purified with a PCR purification kit (Qiagen). The PCR product was phosphorylated with T4 Polynucleotide Kinase, ligated with T4 DNA ligase, and incubated with DpnI (all from New England Biolabs). Following purification with a PCR purification kit (Qiagen), the product was electroporated into MegaX DH10B Electrocomp Cells (ThermoFisher) using a Gene Pulser Electroporation System (Bio-Rad). Dilutions of the library were plated on LB-Ampicillin plates to determine transformation efficiency. The mutant plasmid pool was next subcloned into a promoterless dsRed-Express-derivative plasmid containing a complementary Bxb1 attB sequence for integration into landing pad cells (fully described in Matreyek et al, 2017).^32^ This plasmid was modified by Gibson assembly to contain wildtype *SCN5A*:IRES:mCherry. The mutant plasmid pool was subcloned into the promoterless *SCN5A*:IRES:mCherry plasmid by restriction digest with AatII and AflII (New England Biolabs). A random 18-mer barcode was cloned into the resulting plasmid by restriction digest with Pfl23II (BsiWI) and XbaI (ThermoFisher). Further descriptions of the cloning scheme, barcoding design, and the process to determine which barcode corresponded to which mutation are presented in the supplemental methods.

### Optimization of *SCN5A* triple drug assay

In order to optimize a negative selection SCN5A assay, we used transiently transfected HEK293T cells. All HEK293T cells were grown to 30-60% confluency before transfection, and incubated at 37°C in DMEM supplemented with 10% FBS, 1% penicillin and streptomycin, and 5% CO_2_. Stock solutions were made of 25 mg/ml ouabain octahydrate (Sigma-Aldrich) in hot water, 50 mM veratridine (Sigma-Aldrich) in ethanol, and 100 µg/ml brevetoxin A (PbTx-1; BOC Sciences) in DMSO. Stock chemicals were combined and diluted to the desired final concentration in HEK293T media. Preliminary experiments presented below defined the “triple drug” treatment which unless otherwise specified refers to 5 hours treatment of 10 µM ouabain, 25 µM veratridine, and 250 ng/ml brevetoxin. To test the drug assay performance in transient expression, HEK293T cells were transfected with wildtype or mutant *SCN5A* pIRES2-EGFP plasmids (Clontech) using FuGENE 6 (Promega) on day 0. On day 1, cells were passaged into multiple replicates in 6-or 12-well plates. On day 2, drugs were added for 5 hours after which media changes were performed to remove the drugs. On day 3, dead cells were non-adherent and were removed by multiple media changes, and GFP fluorescence was assessed by analytical flow cytometry. Flow cytometry data were collected on a BD LSR Fortessa with a 488 nm laser and 525/50 nm filter for GFP. For each sample, single cells were identified by Forward and Side Scatter measurements. Highly GFP+ cells were identified, defined as cells with GFP fluorescence greater than 100X the mean background fluorescence of untransfected control cells. These values were used to calculate a normalized fraction of GFP+ cells, defined as:

Normalized GFP+ of sample X = (% cells high GFP+ in sample X) / (Average % cells high GFP+ in untreated control samples). For the triple drug transient expression experiments, three replicates were analyzed for each experimental condition. Experimental conditions were compared with a two-tailed t test, implemented in R.

### *SCN5A* Deep Mutational Scan

The Deep Mutational Scan (DMS) was performed in HEK293T cells stably expressing a mixture of *SCN5A* mutants. For stable expression of this mutant *SCN5A* library, we used a previously described cell line containing a “landing pad” recombination site in the AAVS1 safe harbor locus.^32^ The landing pad consisted of a Bxb1 recombination site adjacent to Blue Fluorescent Protein under a Tetracycline-controlled promoter (HEK 293T TetBxb1BFP). On day 0, a plasmid expressing Bxb1 recombinase (pCAG–NLS–HA– Bxb1; Addgene #51271, a gift from Pawel Pelczar ^50^) was transiently transfected with FuGENE 6 into the HEK TetBxb1BFP line.^32^ On day 1, the promoter-less *SCN5A*:IRES:mCherry plasmid library was transfected with FuGENE 6. To induce expression in the stably transfected HEK293T cells, 1 µg/ml doxycycline was added at least 48 hours before further experiments. After 5 days, cells were FACS sorted to select for mCherry+, BFP-cells, indicating successful integration of the plasmid. FACS was performed with a BD Aria III with 535 nm (excitation) 610/20 nm (emission) and 405 nm (excitation) and 450/50 nm (emission) for mCherry and BFP, respectively. After cell recovery, the triple drug combination was applied to the *SCN5A* library-expressing cells as described above and dead cells were removed by repeated media changes. Pre- and post-drug samples were collected for sequencing, and were pelleted at 300xg for 5 min and frozen at −80° C. DNA was isolated using a Qiagen DNeasy blood and tissue kit. PCR was performed on 1-3 µL of the extracted DNA solution using primers ag445 and ag363, with different 6 bp indices for each replicate sample. Six independently transfected and drug-treated replicates were analyzed. See Table S3 for a list of primers used in the study. Libraries were cleaned with AmpureXP beads (Beckman Coulter) following the manufacturer’s instructions. Quality control of sequencing libraries was assessed by Bioanalyzer (Agilent) and qPCR. Libraries were sequenced on an Illumina MiSeq instrument.

### Calculation of DMS scores

Illumina reads were processed in python to extract barcodes and further statistical analyses were performed in R. Barcode counts were normalized to the overall number of reads in each sample and pooled for each variant. For each variant, a raw DMS score was calculated by taking log_10_(post-drug abundance/initial library abundance). These scores were then averaged across the 6 replicates and normalized using a linear transformation so that the mean nonsense score was 0 and the mean synonymous score was 100. Four nonsynonymous variants absent from the initial plasmid library were excluded. For each variant, mean and standard errors of the DMS scores were calculated from the 6 replicate samples. These standard errors were used to define a 95% confidence interval (1.96xSE). The range of synonymous scores was used to define a wildtype interval. Variants with a score and 95% confidence interval lying completely inside, above, or below, the wildtype interval were classified as wildtype, gain of function, or loss of function, respectively. Variants with a score outside the wildtype range but a 95% confidence interval overlapping the wildtype interval were classified as possibly gain or possibly loss of function. Variants with a score inside the wildtype interval but with a 95% confidence interval outside the wildtype interval were classified as possibly wildtype.

### Voltage clamp of *SCN5A* variants

Mutant SCN5A expression plasmids were generated with Quikchange mutagenesis as described above and sequence verified. 48 hours after transient transfection with wildtype or mutant pIRES2-*SCN5A*-EGFP by FuGENE 6, GFP+ cells were selected by microscopy. Sodium currents were recorded using standard protocols as previously described.^51^ Peak and late currents were measured from a 200 ms depolarization from −120 mV to −30 mV. Peak current density was calculated as the peak current (pA) divided by the capacitance (pF). Late current was divided by peak current to determine a late current percentage. 4-7 replicate cells were analyzed per mutation.

## Supporting information

File S1 DMS Scores

## Acknowledgements

This research was funded by F32 HL137385 (AMG), K99 HL135442 (BMK), P50 GM115305 (DR), 1R01GM109110 (DMF), and an American Cancer Society Postdoctoral Fellowship PF-15-221-01 (KAM). The authors thank the Vanderbilt VANTAGE genomics core for high-throughput sequencing.

## Author contributions

AG performed experiments, led the data analysis, and prepared the paper. BK assisted with library generation, structural biology analyses, and data analysis. KM and DF provided important reagents and experimental guidance. TY performed patch clamp experiments. YW and TS assisted with cloning and cell culture experiments. JS assisted with data analysis. DR provided experimental and data analysis guidance and helped prepare the paper. All authors discussed the results and commented on the manuscript.

## Materials and Correspondence

Materials requests can be made to Dan Roden.

## Data Availability

The Deep Mutational Scan dataset generated during the current study is available in File S1. Any additional data or computer code can be made available upon reasonable request to the corresponding author.

