## Supplement for "Deep Mutational Scan of a cardiac sodium channel voltage sensor"

### Supplemental Methods

Design of *SCN5A* plasmid zone system.

We modified a previously published promoterless dsRed-Express-derivative plasmid[1] to insert wildtype *SCN5A*. The most common allele of the common H558R polymorphism (H558) and the most abundant splice variant in the heart (Q1077del) were used. Wildtype *SCN5A* was modified with Quikchange mutagenesis (Agilent) to introduce synonymous mutations that introduced AflII and AatII restriction sites at amino acid positions 1533 and 1785. A "zone" plasmid containing only AA#1533-1785 was also subcloned into the promoterless dsRed-Express-derivative plasmid. A diagram of the zone system is shown in Figure S2.

Adding barcodes to plasmid library.

To create the barcode, primers ag289 (spacer-BsiWI-AflII-polyN-AatII-XbaI-spacer) and ag290 (reverse complement of the 3' end of ag289) were annealed by heating to 95$^{o}$ for 3 minutes and cooling at 10s/1$^{o}$. The primers were extended to make fully double stranded with Klenow polymerase (NEB). The double-stranded barcode was then subcloned by restriction digest with BsiWI and XbaI into the plasmid containing the *SCN5A* library and electroporated into bacteria as described above.

Subassembly for association between barcode and variant.

See Figure S2 for a diagram of the subassembly process. The plasmid pool was digested with AatII and religated with T4 ligase (both New England Biolabs) to bring the barcode near the mutation. The resulting plasmid pool was electroporated, maxiprepped, and gel extracted to obtain intramolecular ligation products as opposed to intermolecular products. PCR with one primer left of the barcode (ag417) and one primer right of the 12 aa mutational target region (ag414) was done to add Illumina flowcell-binding sequence. The subassembly library was sequenced with Illumina MiSeq 75 base paired end sequencing and the sequencing reads were analyzed with python to identify barcodes that were associated for >80% of reads with a single mutation (or multiple mutations). Subassembly identified a single major associated variant for 7,185/7,205 barcodes (Table S1).

**<u>Supplemental Figures, Tables, and Files</u>**

**Figure S1: Triple drug assay is sensitive to treatment time and ouabain concentration.**

**Figure S2: Single codon mutagenesis.**

**Figure S3: Cloning diagram for Deep Mutational Scan.**

**Figure S4: Abundance of individual amino acid substitutions determined by Illumina sequencing of mutagenized *SCN5A* library in the target 12-aa region**

**Figure S5: Abundance of nonsense, synonymous, and nonsynonymous variants in initial library, pre-drug pool, and post-drug pools.**

**Table S1. Mutations present in barcoded library.**

**Table S2. Electrophysiological parameters of newly studied variants**

**Table S3. Primers used in this study.**

**File S1. DMS scores for all variants.**

### Supplemental Figures

#### Figure S1. Triple drug assay is sensitive to treatment time and ouabain concentration.

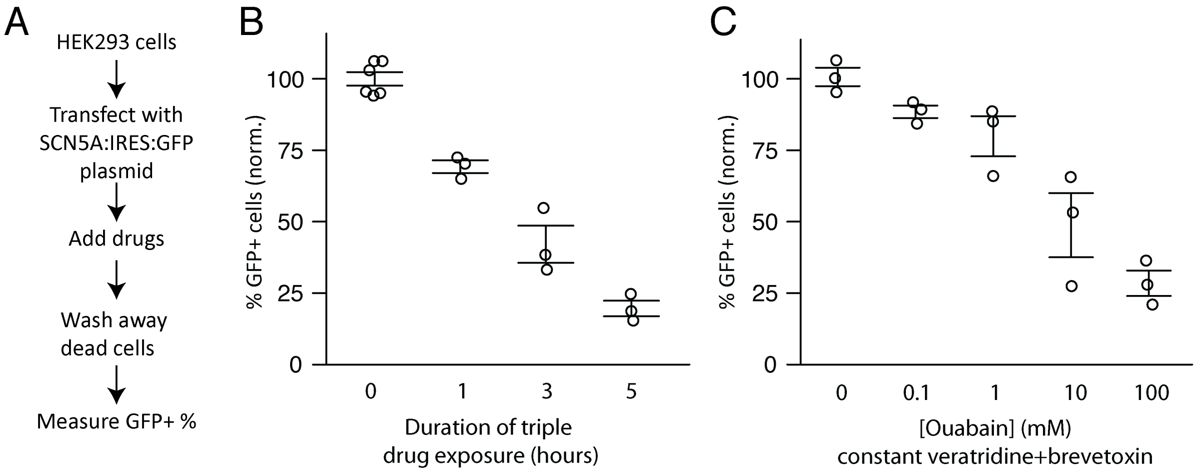


A) Schematic of method to detect selection against *SCN5A* variants after drug treatment. Drug assays were tested as in Figure 2. B) Drug concentrations were kept constant but exposure duration was varied. C) 5 hours treatment with constant veratridine and brevetoxin, while varying the ouabain concentration. Error bars indicate standard error of the mean.

**Figure S2. Single codon mutagenesis.**

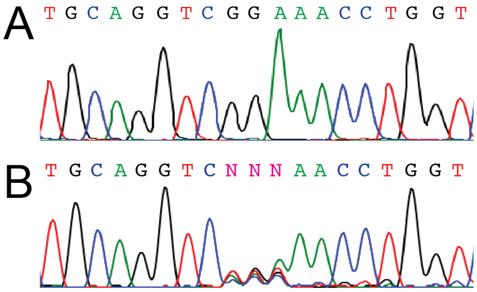


C

| Plasmid | Sequence | Variant |
|---|---|---|
| WT | ...GGTCGGAAACC... | -- |
| G752* pool | ...GGTCNNNAACC... | G752* |
| Clone 1 | ...GGTCACAAACC... | G752T |
| Clone 2 | ...GGTCCCGAACC... | G752P |
| Clone 3 | ...GGTCATGAACC... | G752M |
| Clone 4 | ...GGTCCTAAACC... | G752L |
| Clone 5 | ...GGTCGTCAACC.. | G752V |

A) Wildtype *SCN5A*. B) G752* mutant *SCN5A* library shows a mixture of all possible codons at AA#752. C) Sequencing of individual colonies shows a variety of different mutations at position 752.

**Figure S3. Cloning diagram for Deep Mutational Scan.**

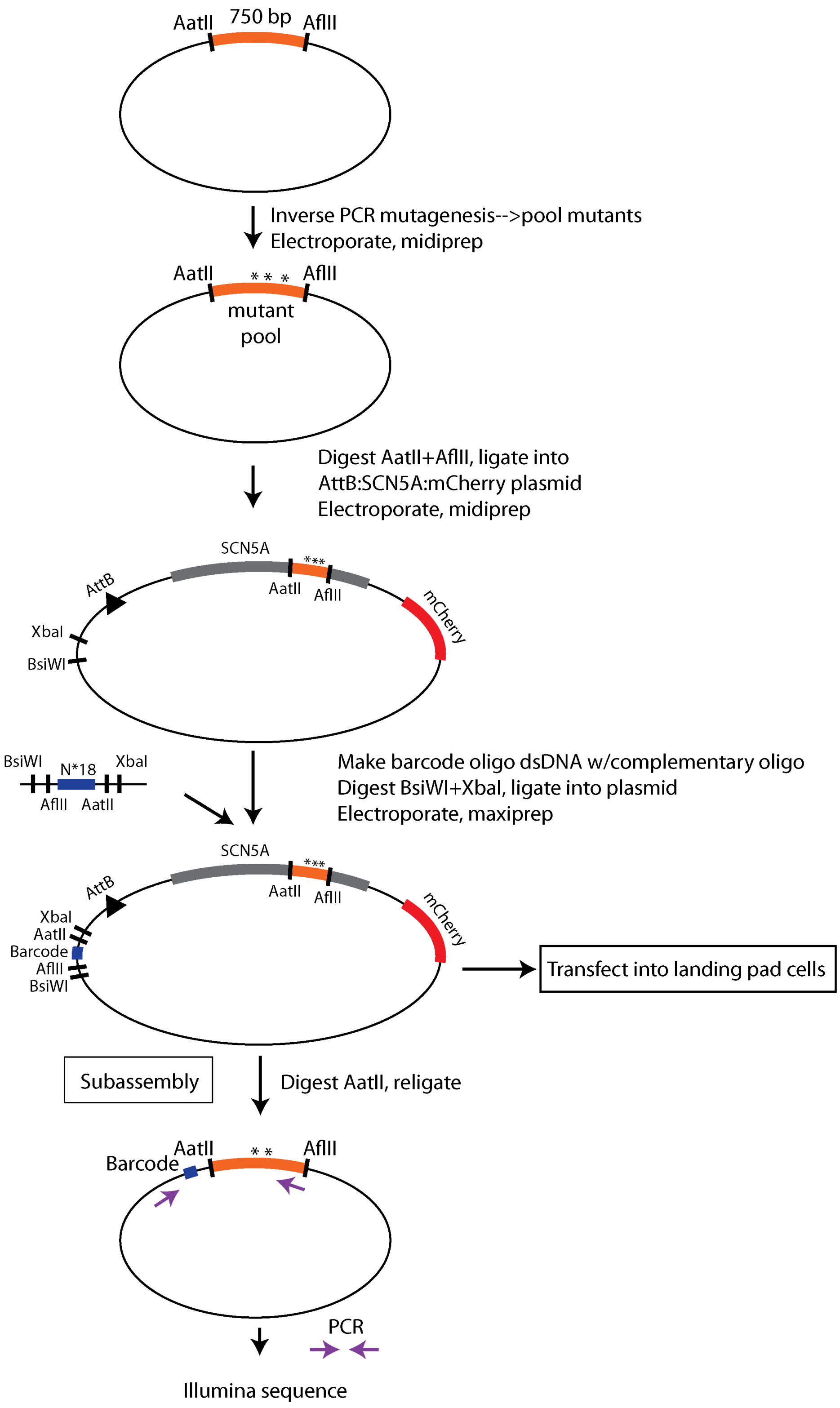

**Figure S4. Abundance of individual amino acid substitutions determined by Illumina sequencing of mutagenized *SCN5A* library in the target 12-aa region.**

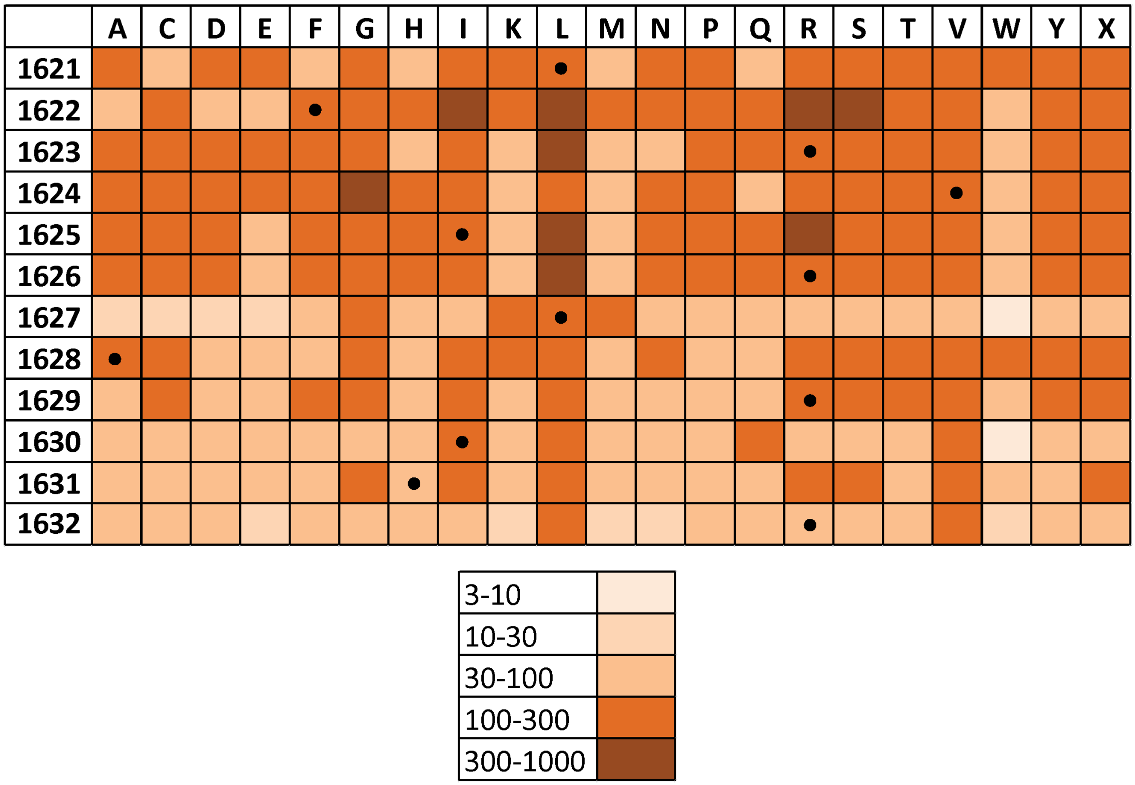


There were a total of 32.7 million reads, and each of the 228 possible single amino acid changes were detected. The number of reads supporting each mutation is shown in the key (in 1000s). The wildtype residues (synonymous variants) are shown with a black dot.

**Figure S5. Abundance of nonsense, synonymous, and nonsynonymous variants in initial library, pre-drug pool, and post-drug pools.**

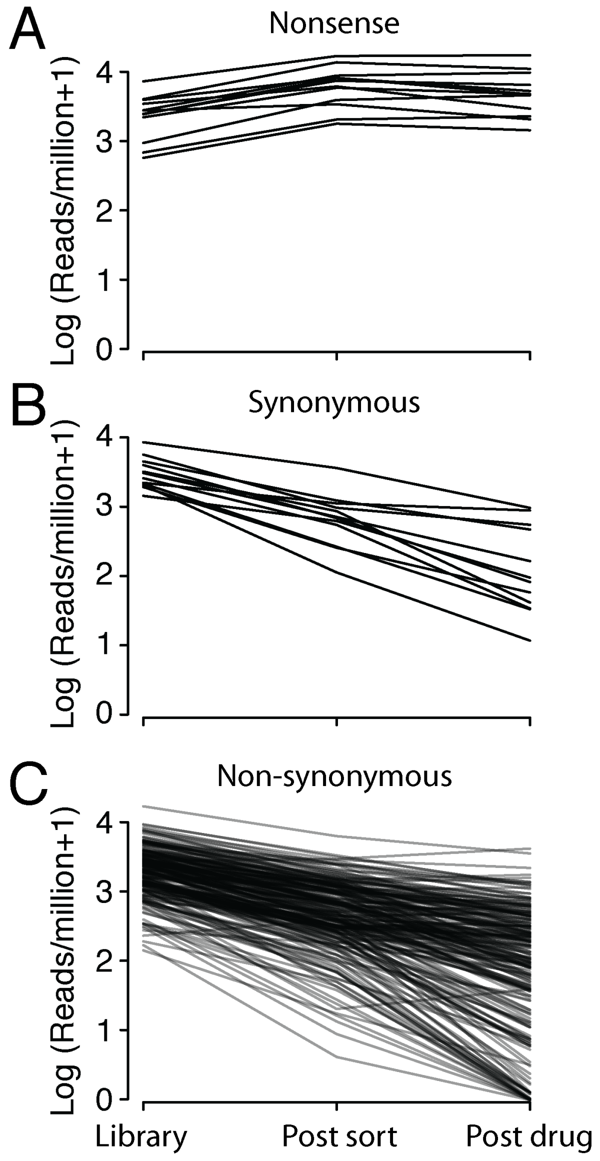


Read counts were normalized to the total number of million reads in each sample and log-transformed.

### Supplemental Tables

#### Table S1. Mutations present in barcoded library

| Mutation Type | # unique barcodes | # unique DNA changes (observed/possible) | # unique protein changes (observed/possible) |
|---|---|---|---|
| Nonsense | 248 | 35/36 | 12/12 |
| Synonymous | 305 | 43/44 | 12/12 |
| Missense | 4277 | 626/676 | 224/228 |
| Wildtype | 166 | 1/1 | 1/1 |
| Multiple | 164 | * | * |
| Indel | 2025 | * | * |
| Conflicting | 20 | * | * |
| Total | 7205 | 705/757 | 249/253 |

### Table S2. Electrophysiological parameters of newly studied variants

| Mutation | # cells | Peak current density (pA/pF) | Normalized peak current density | Late current (% of peak) |
|---|---|---|---|---|
| WT | 7 | 213.3 (17.3) | 100.0% (8.1) | 0.29% (0.07) |
| V1624S | 7 | 163.3 (14.5) | 76.6% (6.8) | 1.95% (0.23) |
| I1630L | 7 | 63.5 (9.2) | 29.8% (4.3) | * |
| V1624E | 4 | 0.0 (0.0) | 0.0% (0.0) | * |

Mean (Standard error).

**Table S3. Primers used in this study**

| Name | Description | Sequence |
|---|---|---|
| ag289 | Barcode forward | attaCGTACGcttaagNNNNNNNNNNNNNNNNNgacgtccTCTAGAgcca |
| ag290 | Barcode reverse | TGGCTCTAGAGGACGTC |
| ag414 | Subassembly right of mutation | 5'[N]AATGATACGGCGACCACCGAGATCTACACTCTTTCCCTACACGACGC TCTTCCGATCTGGCCCCTCGGATCAGTCT |
| ag417 | Subassembly left of barcode | 5'CAAGCAGAAGACGGCATACGAGATTGACCAGTGACTGGAGTTCAGAC GTGTGCTCTTCCGATCTCGGCAATTCCGGACGTACG |
| ag445 | Library seq. left of barcode | 5'[N]AATGATACGGCGACCACCGAGATCTACACTCTTTCCCTACACGACGC TCTTCCGATCTCGGCAATTCCGGACGTACG |
| ag363 | Library seq. right of AttP in genome | 5'CAAGCAGAAGACGGCATACGAGAT[index]GTGACTGGAGTTCAGACG TGTGCTCTTCCGATCTCTCTTCGCCCTTAGACACCAT |
| ag207 | AA 1621* F | NNNTTCCGAGTCATCCGCCTGGC |
| ag208 | AA 1622* F | NNNCGAGTCATCCGCCTGGCC |
| ag209 | AA 1623* F | NNNGTCATCCGCCTGGCCCGAAT |
| ag210 | AA 1624* F | NNNATCCGCCTGGCCCGAATAGG |
| ag211 | AA 1625* F | NNNCGCCTGGCCCGAATAGGC |
| ag212 | AA 1626* F | NNNCTGGCCCGAATAGGCCGCAT |
| ag213 | AA 1627* F | NNNGCCCGAATAGGCCGCATCCT |
| ag214 | AA 1628* F | NNNCGAATAGGCCGCATCCTCAGAC |
| ag215 | AA 1629* F | NNNATAGGCCGCATCCTCAGACTGAT |
| ag216 | AA 1630* F | NNNGGCCGCATCCTCAGACTGATC |
| ag217 | AA 1631* F | NNNCGCATCCTCAGACTGATCCGAG |
| ag218 | AA 1632* F | NNNATCCTCAGACTGATCCGAGGGG |
| ag219 | AA 1621* R | CGTCGGGGAGAAGAAGTACTTCT |
| ag220 | AA 1622* R | GAGCGTCGGGGAGAAGAAGTAC |
| ag221 | AA 1623* R | GAAGAGCGTCGGGGAGAAGAAG |
| ag222 | AA 1624* R | TCGGAAGAGCGTCGGGGAGA |
| ag223 | AA 1625* R | GACTCGGAAGAGCGTCGGG |
| ag224 | AA 1626* R | GATGACTCGGAAGAGCGTCGG |
| ag225 | AA 1627* R | GCGGATGACTCGGAAGAGCG |
| ag226 | AA 1628* R | CAGGCGGATGACTCGGAAGAG |
| ag227 | AA 1629* R | GGCCAGGCGGATGACTCG |
| ag228 | AA 1630* R | TCGGGCCAGGCGGATGACT |
| ag229 | AA 1631* R | TATTCGGGCCAGGCGGATGAC |
| ag230 | AA 1632* R | GCCTATTCGGGCCAGGCG |
| ag109 | AA 752* F | NNNAACCTGGTCTTCACAGGGATTTTCAC |
| ag110 | AA 752* R | GACCTGCAGCATCTCCTCGAATT |
| ag145 | G752R Quikchange | GGAGATGCTGCAGGTCAGAAACCTGGTCTTCAC |
| ag128 | R620H Quikchange | AAGCCACCTCCTCCACCCTGTGATGCTAG |
| ag132 | N1325S Quikchange | GGCATGAGGGTGGTGGTCAGTGCCCTGGTG |
| ag068 | W822X Quikchange | GCTGGCCAAATCATGACCCACCCTGAACACA |
| ag501 | V1624E Quikchange | GACGCTCTTCCGAGAAATCCGCCTGGCCCG |
| ag549 | V1624S Quikchange | CCGACGCTCTTCCGATCTATCCGCCTGGCCCGA |
| ag554 | I1630L Quikchange | CATCCGCCTGGCCCGACTTGGCCGCATC |

**Table S3. Primers used in this study (continued)**

*[N] indicates that either 0, 2 or 4 N's were added to the beginning of the primer and mixed in equal ratios to ensure library diversity upon Illumina sequencing. [index] indicates that multiple 6 base indexes were used for multiplexing multiple Illumina samples.
